# *Host_microbe_PPI* - R package to analyse intra-species and inter-species protein-protein interactions in the model plant *Arabidopsis thaliana*

**DOI:** 10.1101/551275

**Authors:** Thomas Nussbaumer

## Abstract

Intra-species protein-protein interactions (PPI) provide valuable information about the systemic response of a model species when facing either abiotic and biotic stress conditions. Inter-species PPI can otherwise offer insights into how microbes interact with its host and can provide clues how early infection mechanism takes place. To understand these processes in a more comprehensive way and to compare it with experimental outcomes from *omics* studies, we require additional methods to analyse and visualize PPI data. We demonstrate the user-interface *host_microbe_PPI* that is implemented in R Shiny. It allows for interactively analysing inter-species and intra-species datasets from various published *Arabidopsis thaliana* datasets. It enables among other features comparisons of the centrality measurements (degree, betweenness and closeness) and analysis the existence of orthologous proteins in closely related genomes, *e.g.* when gene loss in host and non-host plants is compared. Arabidopsis was used even so the tool can be also applied in other host-microbe systems.

## Introduction

The model plant *Arabidopsis thaliana* was the first plant to have its genome fully assembled in 2000 [1] while in 2009, the interactome, which is represented by pairwise intra-species protein-protein interactions (PPI) became available [2] describing 6,200 interspecies PPI from 2,600 proteins. In the upcoming years, more PPI datasets appeared that can be grouped into biochemical, biophysical, genetic and computational methods [3]. Yeast-two-hybrid (Y2H) techniques, which belong to genetic methods, allowed to systematically describe inter-species PPI between *A. thaliana* to fungi, bacteria and oomycetes such as *Hyaloperonospora arabidopsidis* [3], *Pseudomonas syringae* [3] and *Golovinomyces orontii* [4]. These microbes have an evolutionary distance of several hundred million years and allowed to demonstrate, that effectors from different kingdoms can still target more common host proteins than expected by chance. These Y2H studies enables us to draw several novel scientific conclusions about inter-species PPI: It could for example show, that host proteins are sometimes targeted by several effectors from one particular microbe indicating that if one effector fails to target a host protein, other effectors might take over [4]. This could also show that target proteins are in general more connected to other proteins of the host species within the intra-species protein-protein network. This can be achieved by using approaches such as the centrality measurements [6] or K-shell decomposition [7]. In addition, several studies have used these datasets and applied them to other closely related species or species that are targeted by specific microbe that lakes PPI. Therefore, the ‘interolog approach’ [8] was established which assumes that experimentally verified proteins in one species might improve our understanding of protein-protein interactions in closely related species based on orthologous protein.

In many Y2H studies, computational procedures were used to interpret the inter-species PPI [3,4]. However, in order to make use of the PPI datasets it was necessary to combine different datasets from various different studies. Here, we make use of the R Shiny concept which provides a user interface that allows non-bioinformaticians to execute and analyse PPI data and by combining various heterogeneous datasets for *Arabidopsis thaliana*. Therefore, the R package can be used to directly analyse *A. thaliana* interspecies datasets while it also enables application of methods to other genomes when providing requested input files as described below.

## Material and Methods

### Significance through random selections

The simulation approach enables us to assess whether the amount of distinct host proteins that are targeted by the entire effector repertoire is higher or lower than expected. Therefore, if a total of *N* interactions appear in the interspecies PPI dataset, proteins will be randomly picked using the ‘*sample*’ function in R (parameter: *replace*=TRUE). Within *host_microbe_PPI*, a user can select a value between 10,000-100,000 randomisations to analyse whether the amount of distinct host proteins in the screen is significantly different to that from the simulation (within the 1% or 99% quantile of the results). Customized R scripts are finally used to visualize the observed versus the expected values using a histogram as was seen in Mukhtar et al. [3] and Wessling et al. [4].

### Orthology approach

Orthologous groups for *A. thaliana* to other genomes were extracted from the Orthologous Matrix (OMA) resource [9] when considering following plant species: monocots: *Aegilops tauschii, Brachypodium distachyon, Zea mays, Oryza sativa, Sorghum bicolor, Triticum aestivum* as well as dicots *Arabidopsis thaliana, Arabidopsis lyrata, Medicago truncatula* and *Solanum lycopersicum*. For the intra-species and inter-species PPI, the relative amount of orthologous proteins per species is shown while the presence-absence matrix is used as input for computing heatmap depicting the correlation between species with help of the R Shiny library d3heatmap [10].

### Centrality measurements

Targeted host proteins are also analysed in terms of three centrality measurements degree, betweenness and closeness that have to be provided by the user and were derived from the Arabidopsis Interactome [2] in our test case. Then, for each target protein it shows list whether they are higher or lower than average including a Wilcoxon Rank Sum test to compare differences between intra-species and inter-species PPI.

## Results

The R Shiny instance *host_microbe_PPI* allows to analyse intra-and inter-species PPI datasets in an interactive way as outlined in **Figure 1**. A user can analyse *Arabidopsis thaliana*, it is possible to use existing input datasets by choosing the option “Preload Arabidopsis data!”. Otherwise, a user has to provide various data files as summarized in **Table 1**. To analyse the interspecies PPI data, a user has to provide datasets that describe the host interactome (intraspecies PPI), host-microbe interactions (interspecies PPI), search space, centrality measurements and orthologous gene assignments to other species e.g. determine whether there is a gene loss in non-host systems (see **Table 1** for further explanations of the input datasets). If all required data files are provided, three main analyses options appear: Simulations of randomly sampled proteins versus the inter-species PPI (*Panel A*) to assess whether there are any differences between host targets compared to the remaining proteins; centrality measurements (*Panel B*), to analyse whether there are any characteristics of the host targets in terms of degree, betweenness and closeness compared to the other proteins and an orthology-based analysis (*Panel C*), to reveal differences between genomes that are real hosts in nature and where it might be of interest to understand whether these proteins are also conserved in the non-hosts.

**Table 1.** Overview of the relevant and optional input parameter that are needed in *host_microbe_PPI*.

| <u>ID</u> | <u>Parameter</u> | <u>Required/</u><br><u>Optional</u> | <u>Description</u> |
| --- | --- | --- | --- |
| **A** | **Inter-species protein-protein interaction data** | *Required* | Protein-protein interactions between the host and the microbe (*e.g.* fungus, bacterium) |
| **B** | **Intra-species protein-protein interaction data** | *Required* | Protein-protein interaction within the host (*e.g.* *Arabidopsis thaliana*) |
| **C** | **Search space** | *Required* | All host proteins, that were used in both, inter-species and intra-species protein-protein data |
| **D** | **Centrality Measurements** | *Optional* | All host proteins of the inter-species protein-protein data with all available information (such as degree, betweenness) |
| **E** | **Gene family assignments** | *Optional* | Assignments of host proteins to other species |

**Figure 1.**
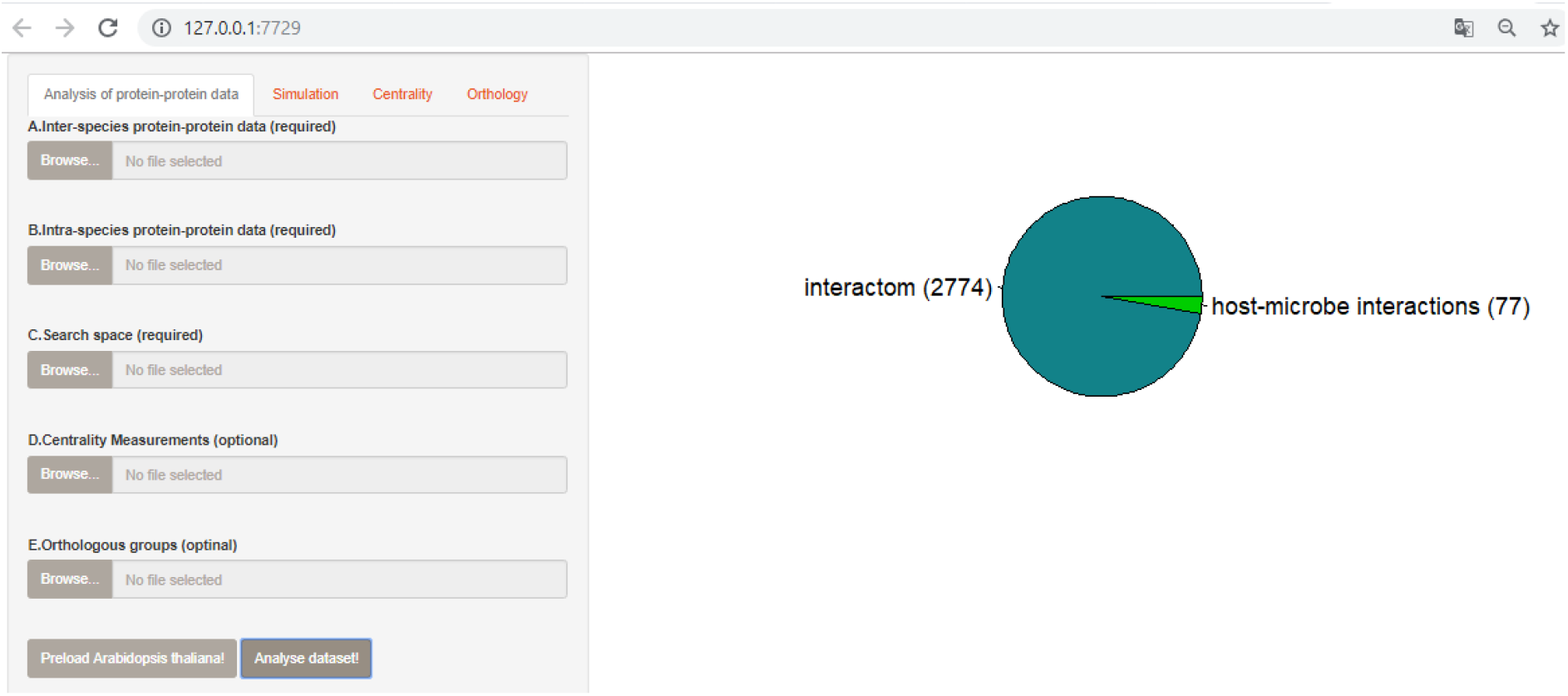
Graphical user interface of host_microbe_PPI. A user can upload the interactome data of the host species (intra-species PPI) containing the protein-protein interactions (PPI), host-microbe interactions (interspecies PPI), search space and optional inputs such as the centrality measurements and the orthologous groups. If all required data is integrated into *host_microbe_PPI*, a user can decide to perform simulations to explore the interspecies PPI, to compare the target proteins in terms of their centrality in the interactome, amount of orthologs between the different genomes as shown in *subfigure A*. *Subfigure B* gives, after all data is loaded, the total amount of proteins that was used the proteins in the interspecies dataset.

### Simulation to analyse whether there are more targets than expected

In the simulation analysis, a user can assess whether the provided inter-species PPI dataset contains more targets than expected when compared to the overall search space (see **Material and Methods**). A user can also select a value between 100-100,000 iterations in the simulation which is adapted from previous studies [3,4]. By default, 10,000 iterations are used when considering the published data from Mukhtar et al. [3] with the host *A. thaliana* and the fungal *Hyaloperonospora arabidopsidis*. For both interspecies PPI, as shown in the original studies, there are far less host targets that are interacting with a microbial effector than expected. Furthermore, as shown in Wessling et al. [4], one may want to compare whether host targets reoccur in different microbes. We used the datasets from *Hyaloperonospora arabidopsidis* and included the data from *Golovinomyces orontii* (**Figure 2**).

**Figure 2.**
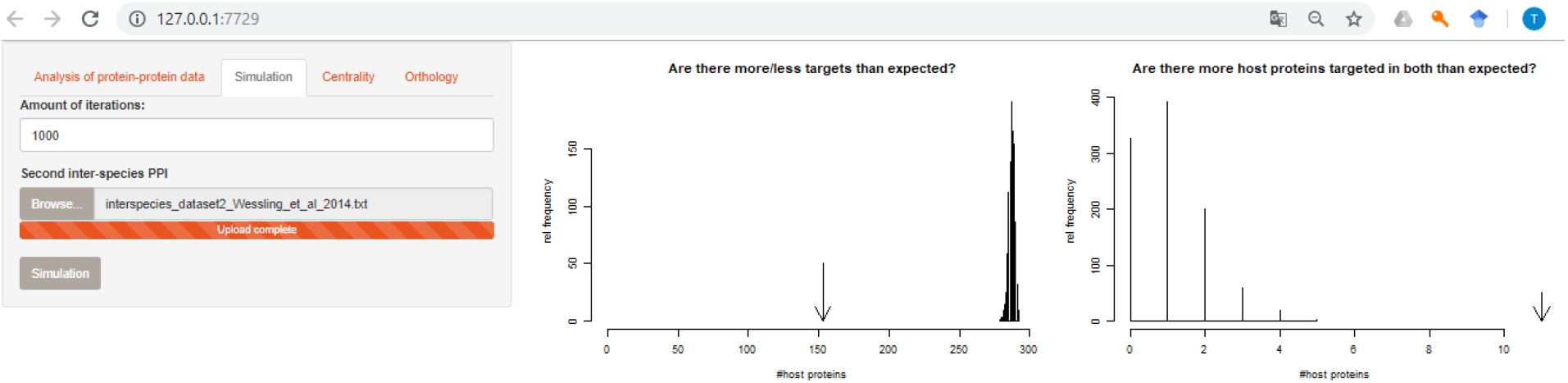
Target simulations. Simulation to analyse whether there are more host targets in interspecies PPI dataset compared to the simulation by randomly picking host proteins from the search space using data from Mukhtar et al. [3] and Wessling et al. [4]. *Subfigure A* shows the amount of interactions can be specified, in *subfigure B* the simulation as histogram is compared with the actual number of distinct host proteins while *subfigure C* shows the common distinct proteins to other screens compared to the simulation.

### Centrality measurement to analyse protein interactions

It is also possible to analyse for differences between intra-species and inter-species datasets in terms of the general centrality measurements such as degree, betweenness or closeness. Assumingly, effectors target rather hub genes, in most cases it is expected that the more strongly connected proteins are preferentially targeted. To prove that, we added a statistical test to compare whether the values of the centrality measurements are significantly different. This was done for the interspecies dataset of *Hyaloperonospora arabidopsidis* (**Figure 3**). The strongest differences were found for all but most strongly for degree (p=1.42e^-23^), btw (p=9.72e^-18^), closeness (p=1.78e^-14^). For *A. thaliana* we have extracted the centrality measurements such as betweenness, closeness and degree were extracted.

**Figure 3.**
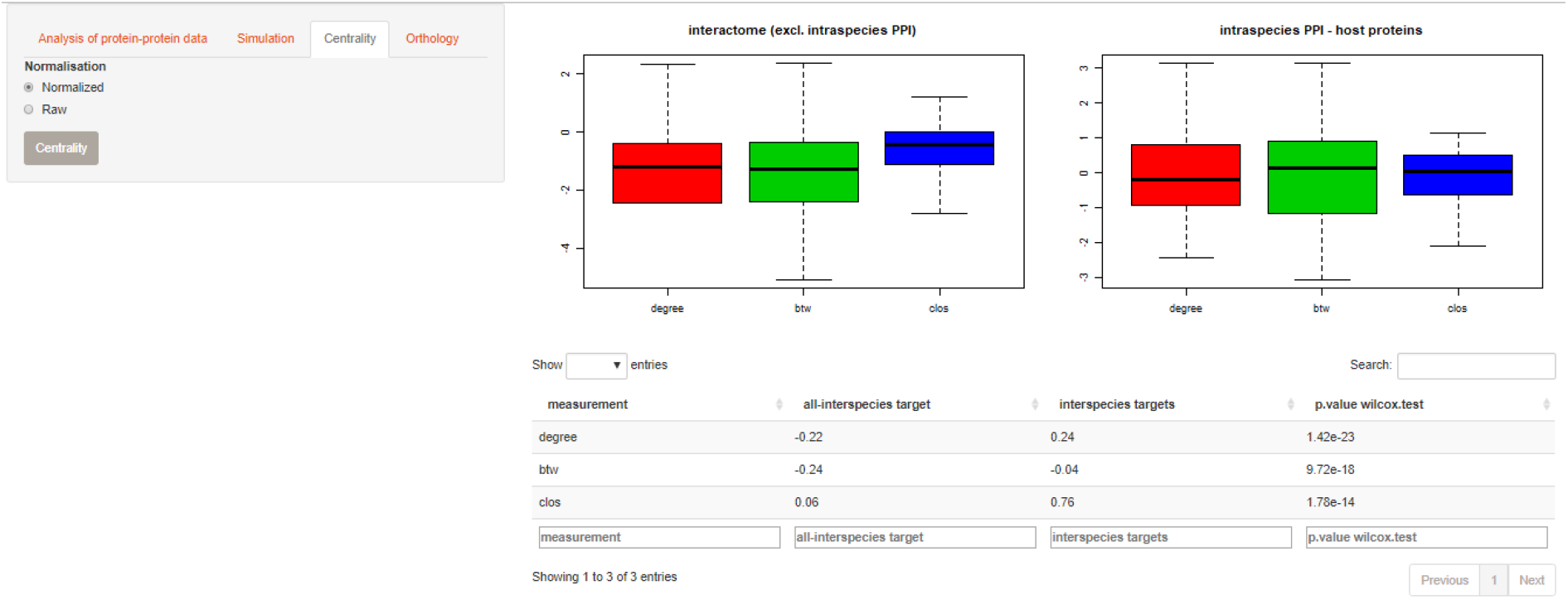
Centrality measurement analysis. Analysis of the comparison of network measurements such as degree, betweenness (btw) and closeness (cls) (*subfigure B*) while the statistical differences between intra-species and inter-species PPI are summarized in an own table (*subfigure C*). Values can be taken as raw values or when Z-transformed (*subfigure A*).

### Orthology-based analysis to analyse host systems versus non-host systems

In particular cases, it might be important to analyse whether host targets are still present in other host genomes or even more interestingly in other non-host genomes. Therefore, **Figure 4a** shows the relative amount of orthologous proteins for the inter-and intra-species PPI dataset. It also shows the correlation between the species based on the presence-absence of single proteins that participate in any interaction (**Figure 4b**). Furthermore, colours depict whether the amount of proteins is significantly higher as expected (**Figure 4b**).

**Figure 4.**
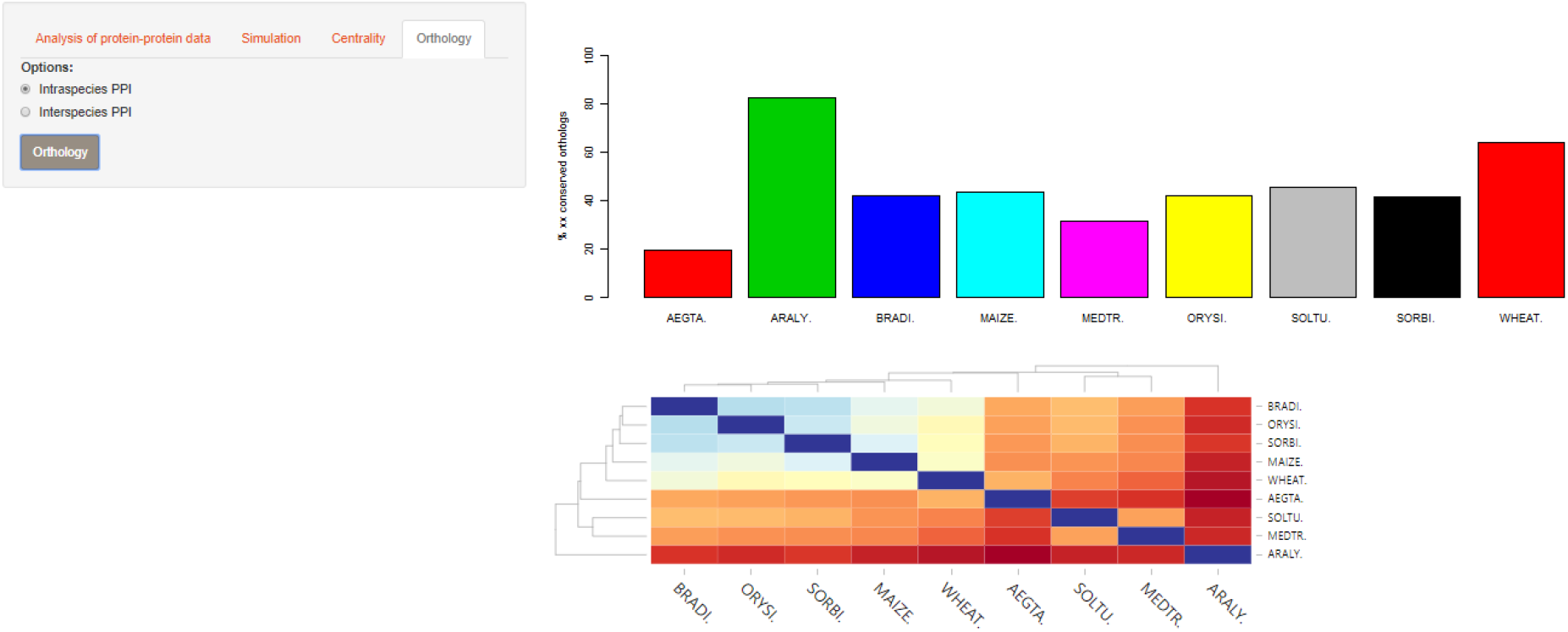
Orthology analysis. Analysis whether the host targets in the interspecies PPI have more or less orthologous proteins in other genomes by depicting the distribution of orthologous genes (*subfigure B*) but also the correlation between the various genomes (*subfigure C*) while *subfigure A* allows to either choose intra-or interspecies PPI.

## Conclusion

With reductions in the sequencing costs and the sequencing of dual RNA-seq datasets becoming more cost efficient more and more datasets become available including datasets containing inter-species PPI. These datasets can help to better understand the PPI data that are already available for selected model species such as *A. thaliana*. *Host_microbe_PPI* combines several analysis methods for intra-and inter-species PPI datasets and can be used to better understand the specific differences of the inter-species PPI in terms of orthology and network centrality.

## Availability

The software is available under https://github.com/nthomasCUBE/host_microbe_PPI. It is also available at ShinyIO under https://nthomascube.shinyapps.io/host_microbe_ppi.

## Acknowledgments

Dr. Alexander Platzer from the Medical University of Vienna is acknowledged for critical reading of the manuscript and for discussion the functionalities. Dr. Jimmy Omony from the PGSB institute at Helmholtz Zentrum Munich is acknowledged for critical reading of the manuscript. Tool is inspired by work within several PPI projects and work with various collaboration partners: Prof. Patrick Schäfer, Rory Osborne, Dr. Jens Steinbrenner, Prof. Paul Birch as well as Stefan Altmann and Prof. Pascal Falter-Braun.

